# Curated high-quality genomes of 39 diverse halophilic archaea

**DOI:** 10.1101/2025.09.09.674397

**Authors:** Zaloa Aguirre-Sourrouille, Jolanda K. Brons, Hanna M. Oksanen, Tessa E. F. Quax, Thomas Hackl

## Abstract

Archaea are widespread and ecologically important microorganisms, yet our understanding of their physiology and evolution is constrained by the limited number of complete genome assemblies available. Haloarchaea have emerged as model organisms for archaeal cell biology, virus-host interactions, and biotechnology. Despite their prominence in hypersaline environments and their potential for industrial applications, high-quality reference genomes remain scarce. Here, we present chromosome-level assemblies for 39 cultivable haloarchaeal strains for which no complete genomes were previously available. Using Oxford Nanopore sequencing, we obtained near-complete assemblies, with 38 strains resolving into single closed chromosomes and additional replicons such as plasmids captured largely in full. These genomes expand the available genomic resources for five haloarchaeal genera and provide a framework for comparative analyses of archaeal metabolism, genome organization, and mobile genetic elements. Given that many of these strains are natural hosts to diverse archaeal viruses, the genomes also represent a critical resource for advancing studies of virus-host interactions in archaea. Beyond fundamental insights into archaeal cell biology and evolution, this dataset will support the development of haloarchaeal model systems and facilitate the exploration of their biomolecules for biotechnological applications.

## Background and Summary

Archaea are intriguing microorganisms that can colonize diverse environments, including extreme habitats such as hypersaline lakes, hydrothermal vents and hot springs. In addition, archaea are found in mesophilic ecosystems, such as oceans and soils, and are part of the human microbiome^1–4^. They play important roles in many ecosystems due to their unique metabolic pathways essential for biochemical cycles^5^. Detection of archaea by metagenome sequencing in many environments shows that they are abundant in the environment^6–8^, and to date, only a tiny fraction of all these species have been cultivated^9^. Archaea have recently also come into focus for understanding eukaryotic evolution. They are phylogenetically closely related to eukaryotes, and the eukaryotic ancestor was likely an archaeon^10–13^. Thus, their prevalence, unique growth conditions and close relation with eukaryotes have contributed to a growing interest in their cell biology, genome organization and metabolism^14^.

Our understanding of archaeal physiology, ecology, and evolution, however, is currently limited by the low number of complete archaeal reference genome assemblies available^15,16^. Only 952 archaeal assemblies on the NCBI genome database are classified as chromosome-level or complete, while for bacteria, there are 80,587 assemblies of this quality level (accessed Sep 2025). The low number of complete archaeal genomes is partly due to the low number of cultured archaeal strains in laboratories. Nevertheless, with the advances in DNA sequencing technology and computational methodologies such as metagenomics, high-throughput sequencing and single-cell genomics, uncultivated microorganisms can be analyzed to investigate microbial communities^17–20^. These technologies have led to the division of archaea into four superphyla: Methanobacteriota, *Nanobdellati and Thermoproteati* (formerly Euryarchaeota, TACK, DPANN, respectively) and Asgard^15,21^.

Within the Methanobacteriota, haloarchaea have emerged as model organisms for studying cell biology, physiology, genetics and virology. Haloarchaea typically inhabit environments with high salt concentrations, such as salt lakes or underground salt deposits^22,23^, ancient halite^24^, table salt^25^, green olives^26^, fish sauce^27^ and mucus within the human gastrointestinal tract^28^. In these environments, which are dominated by haloarchaea, the species diversity is usually low, although cell densities are high. Members of the haloarchaeal genera *Halobacterium, Haloferax*, and *Haloarcula* have become popular model organisms due to their relatively fast doubling times, ability to grow aerobically on defined media and suitability for light and fluorescent microscopy^22,29^.

Halophilic archaea have also been shown to be hosts to abundant archaeal viruses^30–32^. Archaeal viruses, in general, are characterized by a high morphological and mechanistic diversity^33–35^ and are, like their hosts, abundant in diverse environments^7,36–38^. For halophilic archaea specifically, more than 100 viruses have been isolated, which is considerably more than for other archaeal groups^30–32^, rendering this group also an ideal system for studying archaeal host-virus interactions.

Moreover, in recent years, there has been an increasing interest in studying haloarchaea due to their potential for biotechnology. The biomolecules they produce remain active and are stable under high salt concentrations, making them useful for various industrial applications. For example, their enzymes can be used for detergents and bioremediation^39–41^. The polyhydroxyalkanoates and exopolysaccharides they produce could be used to replace commercial non-degradable plastics and polymers^42^. Few haloarchaeal species produce gas vesicles (gas-filled proteinaceous nanocompartments) that can be used to create novel drug delivery systems^43^. Thus, studying haloarchaea will also contribute to harnessing their biotechnological potential^44^.

Here, we present chromosome-level genomes of 39 cultivable haloarchaeal strains for which no reference genome is available (Figure 1). The genomes were sequenced on an Oxford Nanopore MinION sequencer, which turned out to be a highly suited approach for the *de novo* assembly of archaeal genomes. We obtained near-ideal assembly results, with 38 genomes assembling into a single closed main chromosome, in some cases after minor manual curation, and with one genome assembling into a handful of large scaffolds. Additional expected cellular replicons, such as plasmids and mobile genetic elements, were also mostly captured as complete circular sequences.

**Figure 1.**
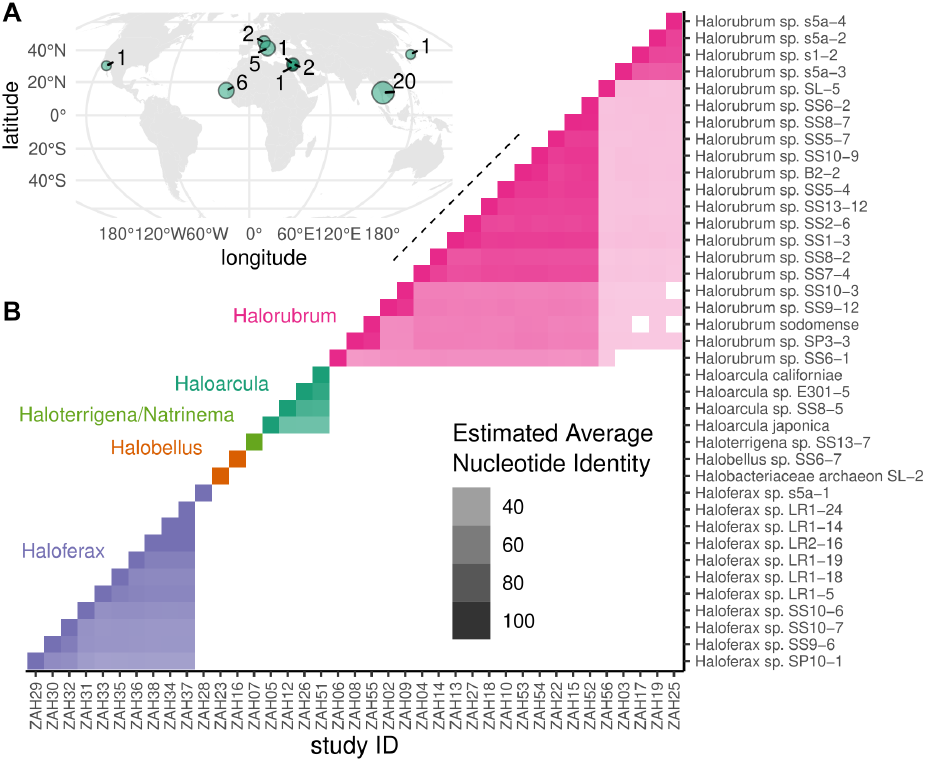
Dataset overview. **A)** Sampling sites across the northern hemisphere. Numbers and circle sizes correspond to the number of genomes per site. **B)** All-vs-all comparison of approximate whole-genome average nucleotide identities (ANI). Strain names are shown on the y-axis and the corresponding internal identifiers on the x-axis. The strains are ordered according to a hierarchical clustering of ANI distances. Genera are shown by colors, and different ANI percentages are encoded by color saturation. The observed grouping of the strains by ANI reflects the classification into five different genera.

The high-quality haloarchaeal genomes presented here will provide a valuable resource for studying haloarchaeal cell biology, metabolism, and evolution. It supports the development of systems for genetic manipulation for these model organisms. Moreover, with most of these strains being known hosts to one or multiple isolated haloarchaeal viruses^30,31,45^, their genomes will also be instrumental for research on archaeal virus-host interactions. Finally, we expect these new genomes of this currently highly underexplored group of microorganisms to advance our understanding of life in extreme conditions and support the development of haloarchaeal biomolecules for biotechnological applications.

## Material and Methods

### Archaeal strains and growth conditions

All haloarchaeal strains (Table S1) were grown aerobically at 42° C in a modified growth medium (MGM) as described previously^46^, if not otherwise stated. *Halorubrum* sp. SS2-6 was grown in YPC^46^, and *Halorubrum* sp. s5a-4 was grown in MGM supplemented with casamino acids.

### Preparation and DNA isolation of archaea

The archaea samples were prepared in accordance with the “Preparation gram-negative and some gram-positive bacterial samples” protocol outlined in the Qiagen genomic DNA handbook, with some modifications. Initially, 4.0 ml of an archaea culture (OD_600_ 0.5 – 0.7) was centrifuged at 5000 x g for 10 minutes at room temperature. Subsequently, the supernatant was discarded, and the cells were washed with 1 ml of phosphate-buffered saline (PBS, Fisher Scientific, Hampton, USA). Following this, the mixture was centrifuged at 5000 x g for 10 minutes at room temperature, and the resulting supernatant was discarded. For the resuspension of the cells, 0.2 mg/mL RNase (Qiagen, Hilden, Germany) was combined with 1 ml of B1 buffer (Genomic DNA Buffer Set, Qiagen, Hilden, Germany). Then, 45 µl of proteinase K (>600 mAU/ml, Qiagen, Hilden Germany) was added to the mixture, which was then incubated at 37°C for 1 hour. After the incubation period, 350 µl of buffer B2 (Genomic DNA Buffer Set, Qiagen, Hilden, Germany) was introduced and mixed by inversion, followed by incubation at 50°C for 30 minutes.

The archaeal DNA was isolated following the “Isolation of genomic DNA from blood, cultured cells, tissue, yeast, or bacteria using genomic-tips” protocol from the Qiagen genomic DNA handbook, with certain adjustments. Initially, a Qiagen genomic tip 20/G (Qiagen genomic-tip 20/G kit, Qiagen, Hilden, Germany) was equilibrated with 1 ml of QBT buffer (Genomic DNA Buffer Set, Qiagen, Hilden), and allowed to empty via gravity flow. After vortexing the sample for 10 seconds at maximum speed, it was subsequently applied to the prepared Qiagen genomic tip 20/G. After passing through the genomic tip, the tip was subjected to three times 1 ml washes using QC buffer (Genomic DNA Buffer Set, Qiagen, Hilden, Germany). Following this, genomic DNA was eluted with 2 ml of pre-warmed (50°C) QF buffer (Genomic DNA Buffer Set, Qiagen, Hilden, Germany) and precipitated by adding 0.7 volumes (1.4 ml) of room-temperature (15–25°C) isopropanol (Sigma-Aldrich, Saint Louis, USA). The mixture was inverted and incubated for 5 minutes at room temperature, followed by centrifugation at 5000 x g for 15 minutes at 4°C, with subsequent removal of the supernatant. The resulting pellet was washed with 1 ml of cold 70% ethanol (4°C) and centrifuged at 5000 x g for 10 minutes at 4°C. This washing step was repeated three times. Finally, the pellet was dried at 50°C for 10 minutes and resuspended in 100 µl of pre-warmed (50°C) Tris-EDTA (TE) buffer (Sigma-Aldrich, Saint Louis, USA). The DNA was dissolved at 55°C for 1 hour, and the DNA concentrations were measured using a Qubit Flex Fluorometer (Fisher Scientific, Waltham, USA). Additionally, the A260/280 and A260/230 ratios were determined using a Nanodrop (Thermo Fisher Scientific, Waltham, USA). The samples were stored at -20°C prior to sequencing.

### Library preparation and Minion sequencing

The archaea DNA libraries were generated using the SQK-NBD114.24 native barcoding kit 24 V14 (Oxford Nanopore Technologies (ONT), Oxford, United Kingdom), following the manufacturer’s instructions. The initial DNA concentration of each sample was adjusted to a starting concentration of 400 ng/µL based on readings from the Qubit Flex Fluorometer. The library preparation involved a DNA repair and end-preparation, a native barcode ligation, and an adapter ligation and clean-up step. After each step, the DNA concentrations were determined using the Qubit Flex Fluorometer. Sequencing was carried out on an ONT MinION device and with two R10.4.1 flow cells, primed and loaded according to the manufacturer’s instructions. Basecalling and demultiplexing were performed with MinKNOW/Dorado at “super-accurate basecalling, 400 bps”.

### Genome assembly and curation

Genomes were assembled from the sequenced read libraries with Flye^47^. Most read sets assembled directly into a single circular contig for the main chromosome and additional smaller contigs for smaller replicons (putative plasmids/prophages). In a few cases, we generated assemblies at different levels of coverage for optimal results. The assembly of *Haloarcula californiae* (ZAH51) was, moreover, manually curated by pruning a redundant repeat and closing the graph into a single circular main contig. Genome graphs were inspected with Bandage^48^. The resulting assemblies were polished with medaka v1.11.3 ^49^. The main chromosomes were all set to have their starts aligned and co-oriented on their single-copy *rpoB1* gene using seq-circ^50^. The GC-coverage structure of the assembly contig was inspected via plots created in R using the packages tidyverse^51^, ggplot2^52^, patchwork^53^, and googlesheets4^54^. Taxonomic classification of the samples was verified with GTDB-tk^55–58^.

## Data Records

Sequencing data and assemblies were deposited at ENA under PRJEB79315.

## Technical Validation

To ensure high-quality genomes, we assessed the assemblies for structural consistency and contamination using a) GC-coverage plots^59^, b) assembly-graph visualizations^60^, and marker-gene-based completeness estimations^61^. In total, we recovered 36 genomes of high quality based on CheckM2 metrics (Completeness > 90%, contamination < 5%) and three that did not meet these thresholds (Figure 2, Table S2). Manual inspection showed that ZAH08 and ZAH15 both assembled into one complete circular main chromosome and that the appearance of incompleteness here relates to a genuine biological signal or potentially the lack of sensitivity in CheckM2’s marker recognition on these archaeal genomes. Similarly, the contamination in ZAH03 also appears linked to biological signals, not contamination, as the duplicated marker genes all appear on the complete circular main chromosome.

**Figure 2.**
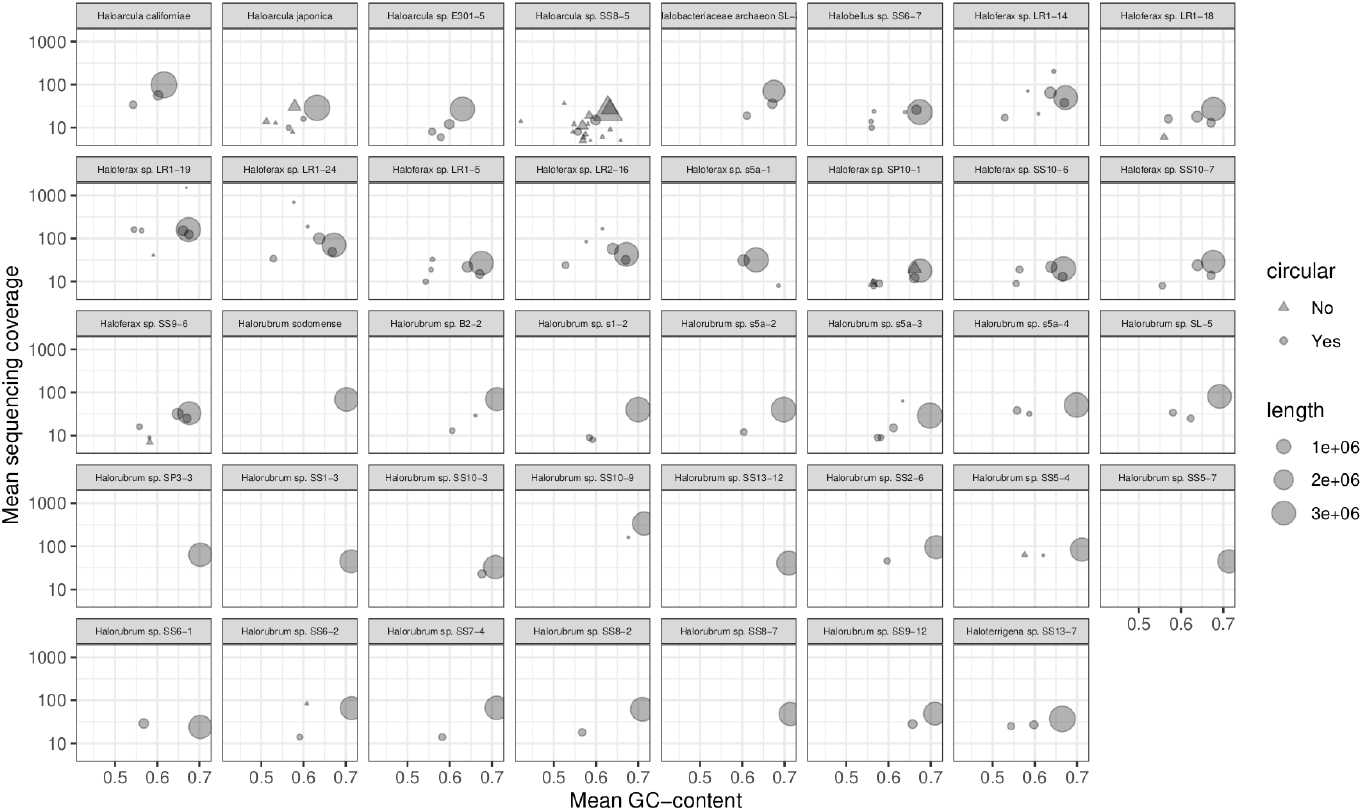
Qualitative genome assembly assessment. Each facet represents the assembly of one strain, with circles and triangles representing assembled circular and non-circular contigs, respectively, and their sizes corresponding to their length in bp. The average sequencing coverage per contig is indicated on the y-axis and the average GC content on the x-axis. The majority of the assemblies comprise a single, circular main chromosome of around 3 Mbp in size, together with zero to four smaller, often circular contigs representing additional replicons in the genomes, such as plasmids.

All our strains have previously been characterized taxonomically based on their partial 16S rRNA gene sequences^30,31,62–64^. Using GTDB-tk^55^ we validated their phylogenetic classification using a whole genome approach (Table S3). We found consistent placement of all 39 genomes between 16S and the whole genome approach. Of note, our *Haloarcula californiae* was classified within the GTDB framework as a member of the species *Haloarcula marismortui*. This is expected, as *H. californiae* and *H. marismortui* form a single species according to GTDB standards. *Haloterrigena* sp. SS13-7 was classified as belonging to the genus *Natrinema*, which is in line with recent findings suggesting a partial merging of the genera *Haloterrigena* and *Natrinema*^*65*^. Halobacteriaceae archaeon SL-2, which initially was not classified at the genus level, was placed into the genus *Halobellum*. Lastly, the six *Halorubrum* isolates SS9-12, s5a-3, SS10-3, s1-2, s5a-2, and s5a-4, could not be classified within GTDB at the species level, suggesting that they represent, to date, undescribed *Halorubrum* species.

## Supplementary Tables

Table S1: Strain and assembly overview

Table S2: Assembly completeness (checkm)

Table S3: Taxonomic classification (GTDBtk)

## Code availability

Utilized software, versions and relevant parameters:

- MinKNOW v23.11.7, Bream v7.8.2, Dorado v7.2.13, MinKNOW Core v5.8.6
- R v4.3.3, tidyverse 2.0.0, ggplot2 v3.5.0, googlesheets4 v1.1.1, patchwork v1.2.0
- Flye v2.9.2-b1786, `‵--nano-hq‵`
- Bandage v0.9.0
- CheckM2 v1.0.2
- Medaka v1.11.3
- seqkit v2.5.1
- minimap2 2.26-r1175
- seq-circ sha:82aed01 (https://github.com/thackl/seq-scripts)
- GTDB-Tk v2.3.2

## Acknowledgments

TEFQ and ZAS were supported by funding from the Hector Fellow Academy. TEFQ was supported by a Vidi grant (Vidi.223.020) from Dutch Research Council (NWO) and an ERC starting grant (ARCVIR, 101039446). HMO was supported by the University of Helsinki and the Research Council of Finland by funding for FINStruct and Instruct Centre FI, part of Biocenter Finland and Instruct-ERIC. We thank the Center for Information Technology of the University of Groningen for their support and for providing access to the Peregrine and Hábrók high-performance computing cluster.

## Author contributions

ZAS, HMO, TEFQ, and TH designed the study. ZAS and JKB performed the laboratory work and sequencing. TH carried out the computational work. All authors contributed to writing the manuscript. TH and TEFQ supervised the project.

